# BAT: an integrated pipeline for gene tree construction, annotation, and functional inference

**DOI:** 10.64898/2026.05.07.721474

**Authors:** Benjamin D. Sheppard, Brian Behnken, Adam Steinbrenner

## Abstract

Gene family functional exploration often requires analyzing motifs, domains, and associated datasets (e.g. gene expression) in the phylogenetic context of a gene tree. As genomic resources become more abundant, local pipelines are needed to analyze gene families of interest with project-specific resources. Here we present BLAST-Align-Tree (BAT), a bioinformatic pipeline for automated gene family phylogeny construction and annotation to enable gene tree exploration. BAT combines a BLAST search of local genome databases with a robust and flexible gene tree construction pipeline that enables multiple modes of annotation. Output visualizations display experimental datasets, custom regex specified amino acid motifs, and protein HMM domain annotations. For flexibility, BAT runs locally and is independent of pre-existing databases, allowing the easy incorporation of custom genomes and datasets. Three primary case studies described here demonstrate the utility of BAT for inferring the function of homologs and orthologs within characterized gene families. BAT is suitable for fine scale phylogenomic analysis of gene families across the tree of life, and default genomes available on installation span model eukaryotes.

## INTRODUCTION

Characterizing genes and their functions is fundamental to biological research. Obtaining and organizing gene coding sequences is a starting point for understanding protein structure and function, and is critical for hypothesis generation prior to experimental work. For example, forward and reverse genetic screens require understanding the nature of candidate genes, as well as any closely homologous genes which could have similar functions. Screens in non-model organisms are especially reliant on careful gene family analysis to compare genes of interest with characterized homologs in genetic model systems (1). Phylogenetic analysis and gene tree construction is also needed to infer orthology (implying potential functional similarity), and for distinguishing orthologs from duplicated genes that existed prior to speciation, i.e. out-paralogs (2). For rapidly evolving gene families or duplication-rich lineages such as plants, pre-computed sets of orthologs with graph- or tree-based methods may also have errors due to the specific clustering, alignment, pruning, and tree construction methods used during gene family construction (3). Finally, at a practical level, molecular biologists also need to inspect homologs for presence of relevant amino acid motifs, protein domains, and other characteristics such as expression patterns.

Gene tree analysis often entails a sequence homology search, alignment of sequences, and phylogenetic analysis (4). The past 40 years have seen major advances in the computational tools available for these tasks (5). BLAST (6), first introduced in 1990, remains the standard for sequence homology searches. Meanwhile, a myriad of well-developed tools are now available for sequence alignment including Clustal Omega, MAFFT, and MUSCLE (7–9), and for tree construction, such as PhyML, RAxML, FastTree, and IQ-TREE (10–14).

Genome-scale orthology methods and databases, including OrthoFinder, OMA, OrthoDB, eggNOG, PLAZA, and GreenPhylDB, (15–20) provide frameworks for inferring orthogroups, hierarchical orthologous groups, gene-family histories, and functional annotations across many species. These resources are valuable for broad comparative genomics, but they are less suited to user-defined exploration of a specific gene family when the relevant genomes, gene models, expression datasets, or feature annotations are local and project-specific.

For tree construction and annotation, existing bioinformatics pipelines can also be complex and lack rapid customization features. Few tools couple local query-driven homolog retrieval, tree construction, and multiple functional annotation modes in a single workflow. For example, NGPhylogeny.fr (21) streamlines tree building from query sequence to inferred gene tree while providing multiple options for alignment and tree construction. Existing visualization tools such as ETE, ggtree, and iTOL allow powerful gene tree annotation, but are not linked to an upstream gene search pipeline (22–24). Instead, data integration requires custom scripting or manual integration. The increasing number of genome annotations, including multiple reference genomes for a single species, further complicates gene tree analysis (25). Therefore, new tools are needed that efficiently combine gene tree construction with putative functional data in a single pipeline.

Here we describe an integrated gene tree construction and annotation pipeline, BLAST-Align-Tree (BAT). BAT is designed as a bottom-up gene family construction pipeline, which complements existing tools. It is a single command line interface pipeline for query-driven homolog retrieval from local databases, flexible alignment and tree construction, and immediate visualization. Additionally, BAT can annotate protein functional domains and amino acid motifs beside tree entries, to facilitate functional hypothesis generation. Flexibility is built into the pipeline with multiple options for sequence alignment (Clustal Omega or MAFFT) and for tree construction (FastTree or RAxML). As outputs, BAT overlays gene trees with both the multiple sequence alignment (including optional user-provided amino acid motifs and protein domains) and a heatmap visualization of user-supplied gene-linked quantitative data. In summary, BAT is a unified workflow for gene tree construction and annotation that enables the rapid and iterative exploration of gene families of interest.

## METHODS

### Console Commands

The installed console command (blast-align-tree) initiates the python script blast_align_tree/cli.py which conducts the BAT Homology Search step (see **Figure 1**) and all steps through the Gene Tree Inference step before invoking the bundled R script visualize_tree.r for visualization. The installed command (bat-genome-selector) invokes a Tkinter graphic user interface that assists with building queries based on the genomes available in the user’s /genomes/ folder. The helper command (blast-align-tree-fetch) downloads additional bundled reference genome FASTAs for BAT. See README at https://github.com/steinbrennerlab/blast-align-tree for additional details for installation and terminal commands.

**Figure 1:**
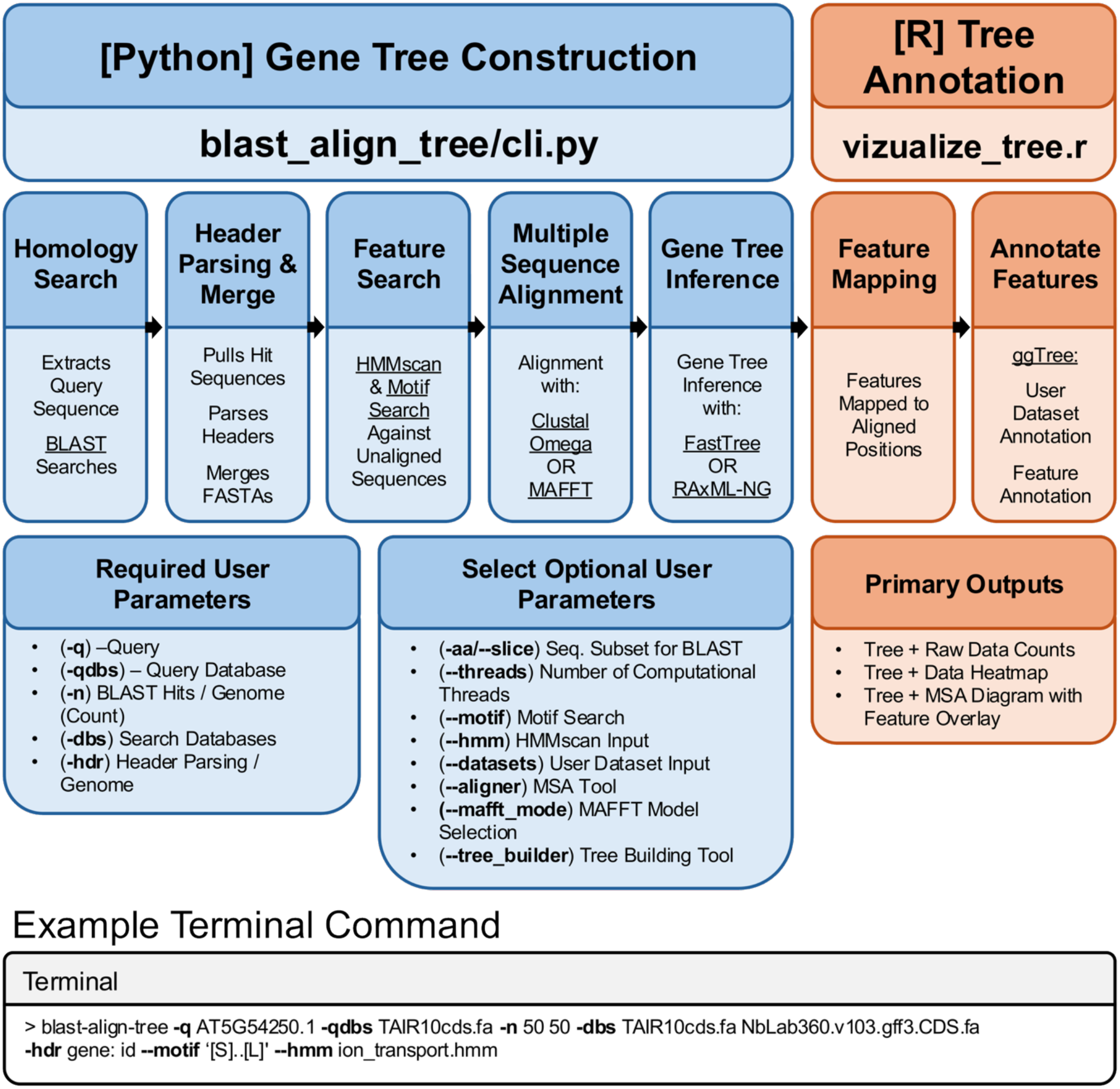
The BLAST-Align-Tree (BAT) pipeline. The top row shows the two major stages of the BAT pipeline. The second row details each step within the pipeline associated with corresponding stages above. The third row shows user-specified input parameters and pipeline outputs. “Example Terminal Command” shows a sample command simplified from the query used to generate **Fig. 3c-e**. Option flags (bold) and key packages (underlined) throughout.

### Homology Search

BAT uses the provided query sequence(s) (-q) to identify homologous sequences from local genome databases via BLAST. The pipeline supports two BLAST search modes: tblastn for nucleotide queries of CDS (Coding Sequence) databases and blastp for protein queries of protein databases. the BLAST mode is selectable via (--blast_type), tblastn is the default. First BAT extracts the query sequence from the specified database (-qdbs), then translates nucleotide sequences to amino acids using the standard genetic code (tblastn mode only). Next BAT conducts BLAST searches in parallel for each provided query sequence against each target genome database (-dbs). The number of retained BLAST hits per database search is limited by (-n). Optionally, the (-aa/--slice) option enables the use of a specified segment of the query sequence for BLAST searching. Selected segments are user specified not identified using BAT (e.g. using InterPro’s online search function (26, 27) to identify the amino acid sequence that codes for a particular protein domain in the query). Note that BLAST hits passed to the Header Parsing and Merge step are always full-length sequences (i.e. hits are not trimmed when (-aa/--slice) is used). The maximum number of parallel searches is specified by the option (--threads), the default is half the maximum. The full alignment report and list of matching sequences is reported for each search.

### Header Parsing and Merge

The Header Parsing and Merge step (see **Figure 1**) extracts all identified homologous sequences, edits sequence headers, and merges the results into a single FASTA file. Hit sequences are extracted from their respective genome databases and processed. For tblastn results (nucleotide sequences), any internal stop codons are removed in-frame to preserve reading integrity, which may occur in pseudogenes or frameshifted sequences. Nucleotide sequences are translated to amino acids, and all sequence headers are standardized using database-specific parsing rules (-hdr). For each database, the (-hdr) parameter specifies a token to search for in the original FASTA header. The parser extracts the text immediately following this token (up to the first whitespace) to use as the sequence identifier. This standardization step allows integration of databases with variable header formats. Note that multiple distinct sequences may receive identical parsed identifiers if a genome encodes multiple splice variants of a gene and (-hdr) specifies extraction of the gene name rather than transcript ID. After header standardization, FASTA files from all databases are merged into a single file. Sequences with identical parsed headers are deduplicated (also possible when using multiple query sequences with overlapping results), and only the first occurrence retained in the analysis.

### Feature Search

The Feature Search step (see **Figure 1**) identifies structural features from the amino acid sequences of identified homologous sequences. Two optional types of protein annotation are available via BAT. The (--motif) option accepts amino acid patterns in either PROSITE or Regex (default) syntax, which are scanned against unaligned sequences to identify functional motifs. The (--hmm) option accepts HMMER3 profile HMM files (.hmm), which are used to detect conserved protein domains using hmmscan (28). Hmmscan gathering thresholds are enabled for stringent filtering. Detected features are recorded with unaligned amino acid coordinates for later mapping onto the multiple sequence alignment by visualize_tree.r.

### Multiple Sequence Alignment

The Multiple Sequence Alignment (MSA) step (see **Figure 1**) aligns the deduplicated protein sequences using one of two alignment tools. The default is Clustal Omega (7). Alternatively, MAFFT (8) is available via the (--aligner) option. Once MAFFT is selected, the (--mafft_mode) parameter can specify the MAFFT alignment model. BAT supports MAFFT: “auto”, “linsi”, “ginsi”, “einsi”, “fftnsi”, “fftnsi-1000”, “fftns2”, “fftns1”, “nwnsi”, “nwns2”, and “parttree” alignment models. If the alignment model is not set by the user, MAFFT selects a strategy automatically. Regardless the MAFFT alignment strategy is logged for pipeline reproducibility, although some models are inherently stochastic. For information about MAFFT model selection and reproducibility see Katoh *et al. 2019* (8) and (https://mafft.cbrc.jp/alignment/software/). Both alignment tools produce a FASTA formatted MSA.

### Gene Tree Inference

The Gene Tree Inference step (see **Figure 1**) builds the tree and calculates support values using the MSA file. The default tree builder is FastTree (13), which generates SH-like local support values but does not calculate bootstrap values. The option (--tree_builder) is used to specify the alternative tree builder RAxML-NG (12) which calculates bootstraps in addition to tree construction. The option (--raxml_seed) allows the user to specify a random seed for construction and bootstrap reproducibility. If an error occurs during tree construction or bootstrap calculation, BAT will try again with fewer threads. The inferred tree is written in Newick format and used directly in downstream visualization steps.

### Feature Mapping

The Feature Mapping step (see **Figure 1**) performed by the bundled R script “visualize_tree.r”, which invokes automatically following Gene Tree Inference. The script collects the MSA FASTA, Tree Newick, feature mapping files, and comma- or semicolon-separated user datasets specified using the (--datasets) flag. Further feature mapping is conducted to convert the un-aligned positions of previously identified sequence features to the aligned MSA file for final visualization.

### Feature Annotation

The Feature Annotation step (see **Figure 1**) uses the R package ggtree (23) to create publication-quality gene tree visualizations. The script generates three visualizations: 1) a gene tree with sequence tips colored by genome origin that also displays raw counts (values) of user provided data (e.g., gene expression, functional annotations) aligned to tree tips; 2) a gene tree with heatmap visualization of user-supplied quantitative data aligned to tree tips; and 3) a gene tree with a visualization of the MSA adjacent to the tree with optional feature overlays (motifs/domains) that appear as colored polygons. Once the initial tree is complete, the R script can be re-run independently using the console command (Rscript “<bundled-visualize_tree.r>”) to adjust visualization parameters (colors, node labels, support value display) or to subset or re-root the tree for publication-quality outputs without repeating the complete pipeline.

### Klepikova Atlas For BAT

To provide a default expression context for the BAT, we constructed a gene-level *Arabidopsis thaliana* expression atlas sourced from Klepikova’s developmental transcriptome atlas data (29). Raw reads were retrieved from NCBI BioProjects PRJNA314076 and PRJNA324514. The SRA accessions used were Dry Seeds (SD.d) (SRR3581731, SRR3581892), Root (R) (SRR3581356, SRR3581836), Internode (IN) (SRR3581705, SRR3581871), Petals of the mature flower (F.PT.ad) (SRR3581688, SRR3581854), Carpels of the mature flower (F.CA.ad) (SRR3581685, SRR3581851), and Leaf, mature (L.lg) (SRR3581681, SRR3581847). Processing of BAMs was performed according to parameters prescribed in Sullivan *et al. 2019* for the BAR eFP Browser (http://bar.utoronto.ca) (30), excepting the use of DESeq2 instead of RPKM for improved normalization in cross-tissue comparison (31). The resulting matrix is distributed with BAT as “klepikova_atlas_subset.txt”. See **Supplementary Table 2** for per-tissue BAT log2 values compared against per-SRR eFP RPKM values transformed as log2(RPKM + 1) including tissue rankings and Pearson correlations for five genes with contrasting expression patterns. Scripts for dataset generation are available at https://github.com/BQBehnken/klepikova_expression_context.

### Model Eukaryote Genome Curation

BAT includes a selection of model eukaryote genomes available via the (blast-align-tree-fetch -- all) command. Coding sequence FASTAs for all species were obtained from their respective repositories and formatted into BLAST-searchable nucleotide databases using makeblastdb (32). “Filename” identifies the file in the BAT repository. “Renamed” indicates the source name if different in the repository beyond trimming of “from genomic”. *Arabidopsis thaliana* (TAIR10 (33) filename TAIR10cds.fa) was obtained from TAIR (arabidopsis.org), *Phaseolus vulgaris* (v1.0 (34) filename Pvul218cds.fa), and *Vigna unguiculata* (v1.1 (35) filename Vung469cds.fa, renamed from download file Vunguiculata_469_v1.1.cds_primaryTranscriptOnly.fa) were obtained from Phytozome 12 (36) (phytozome-next.jgi.doe.gov). *Saccharomyces cerevisiae* (S288Cv64.5.1 (37) filename S288C_scer_cds_canonical.fasta, renamed from download orf_genomic.fasta) was obtained from the Saccharomyces Genome Database (37) (sgd-archive.yeastgenome.org/sequence). *Drosophila melanogaster* (Release 6, ISO1 MT (38) filename D_melanogaster_R6+_ISO1_MT_cds.fna), *Danio rerio* (GRCz12 (39) filename D_rerio_GRCz12tu_cds.fna), and *Caenorhabditis elegans* (WBcel235 (40) filename C_elegans_WBcel235_cds.fna) were obtained from NCBI RefSeq (41) (ncbi.nlm.nih.gov). *Nicotiana benthamiana* genomes (Niben v1.0.1 (42) filename Niben101cds.fa) and (NbLab360.v103 DNA and protein FASTAs (43) filenames NbLab360.v103.gff3.CDS.fasta and NbLab360.v103.gff3.CDS.fasta.AA.fasta) were downloaded from Sol Genomics (44) (https://solgenomics.net). Some genomes obtained from Ensembl Biomart: human (*Homo sapiens*) (GRCh38 (45) filename human_GRCh38_cds_canonical.fa), chimp (*Pan trogolodytes*) (Pan_tro_3.0 (46) filename chimp_NHGRI_mPanTro3-v1.1_cds_canonical.fa), mouse (*Mus musculus*) (GRCm39 (47) filename mouse_GRCm39_cds_canonical.fa), and rat (*Rattus norvegicus*) (mRatBN7.2 (48) filename rat_mRatBN7.2_cds_canonical.fa), exposed problematic headers that prevented isoform compression using (-hdr). For these genomes, canonical transcript FASTAs were obtained from Ensembl Biomart (49) using custom python scripts, that also filtered out alternate loci scaffolds during download. Alternative loci are supplementary sequences in human reference assembly GRCh38 to represent divergent haplotypes at highly variable regions (50) but appear as gene duplicates in the BAT pipeline if unfiltered.

### Case Study Genome Curation

Additional genomes for BAT case studies were obtained, and their databases were built as described above. *Solanum lycopersicum* (ITAG4.0 (51) filename Solyc_ITAG4.0_CDS.fa) was downloaded from Sol Genomics (44) (https://solgenomics.net). *Arabidopsis lyrata* (v2.1 (52) filename Alyrata_384_v2.1.cds_primaryTranscriptOnly.fa), *Brassica rapa* (O_302V_v1.1 (53) filename BrapaO_302V_711_v1.1.cds.fa), *Glycine max* (Wm82.a6.v1(54) filename Gmax_880_Wm82.a6.v1.fa), and *Eutrema salsugineum* (v1.0 (55) filename Esalsugineum_173_v1.0.cds.fa), were downloaded from Phytozome (36) (phytozome-next.jgi.doe.gov). *Cucumis sativus L. var. sativus cv*. Gy14 (Gy14v2 (56) filename Gy14_v2.cds.fa), *Citrullus lanatus* (97102v2.5 (57) filename watermelon_97103_v2.fa), *Lagenaria siceraria* (USVL1VR-Ls (58) filename USVL1VR-Ls_CDS_v1.fa), *Cucumis melo* (DHL92v4 (59) filename CM4.0_transcripts.fasta), *Cucurbita maxima* (Rimu v1.1 (60) filename Cmaxima_v1.1.cds.fa), and *Cucurbita pepo* (MU-CU-16 v4.1 (61) filename Cpepo_v4.1.cds.fa), were downloaded from Cucurbit Genomics (62) (http://www.cucurbitgenomics.org/v2/).

## RESULTS

### BAT is implemented as two interconnected scripts executed sequentially

The console command “blast-align-tree” activates pipeline scripts that parse inputs, execute the BLAST (6) homology search, processes hits, perform the multiple sequence alignment, construct the gene tree, and invoke the R script (**Figure 1**). Once invoked, the R script (visualize_tree.r) generates publication-quality gene tree visualizations incorporating annotations alongside the phylogenetic analysis (**Figure 1**). See Methods for additional details.

### Standard BAT gene trees are consistent with previously documented gene families in plants and animals

To demonstrate typical BAT gene tree outputs we constructed a gene tree for the well described mammalian NAIP (Nucleotide-Binding LRR Apoptosis Inhibitory Protein) gene family. The generated tree recovers single copies of NAIP in both *Homo sapiens* and *Pan troglodytes* and demonstrates the previously documented lineage expansion in mouse (*Mus musculus*, NAIP1, NAIP2, NAIP5, and NAIP6) and rat, (*Rattus norvegicus* NAIP5 and NAIP6) (63, 64) (**Supplemental Figure 1**).

Similarly in plants, a tree built with the *A. thaliana* gene AT5G46330 as BLAST query recapitulates the repeated lineage-specific loss of the plant RLK (Receptor-Like Kinase) immune receptor FLS2 (Flagellin Sensing 2) in Cucurbits (Gourds). Extant Cucurbit FLS2 orthologs form a well-defined clade in the BAT gene tree consistent with the species phylogeny (65). The Gene tree shows the retention of FLS2 in *Citrullus lanatus* (Watermelon), *Lagenaria siceraria* (Bottle Gourd), and *Cucumis sativus* (Cucumber). However, FLS2 is absent in *Cucumis melo* (Melon), *Cucurbita maxima* (Squash), and *Cucurbita pepo* (Zucchini) (**Supplemental Figure 2)**. The loss of FLS2 in many domesticated Melon accessions was previously described (66). We note that drawing conclusions regarding orthology or gene loss is greatly complicated by BLAST parameters, and that the full searches used here analyzed 30-50 hits per search. BAT searches can be easily adjusted to include large numbers of hits (Supplemental Table 1). These test trees demonstrate the utility of the BAT pipeline for recovering complex events in gene families consistent with the literature across plant and animal kingdoms.

### Case studies demonstrate BAT’s data and feature annotations utility

Here we include three case studies detailing advanced applications of the BAT pipeline for incorporating functional data in gene trees.

#### 1. Gene Tree Annotation Using RNA-seq Data

Displaying transcriptomic data alongside a phylogenetic analysis can quickly highlight important patterns. For example, RLK receptors form a large family in plants. However, only certain receptors are involved in immune recognition and signaling (67). Immune receptors are often induced by PAMPs (Pathogen Associated Molecular Patterns), distinguishing them from RLKs with developmental functions (68). Bjornson *et al*. characterized the transcriptome of the plant *A. thaliana* following treatment with various PAMPs (68). Incorporating this transcriptomic dataset into a multispecies RLK gene tree could enable prediction of novel, uncharacterized RLK immune receptors.

The BAT pipeline was used to construct a gene tree targeting RLK sub-family LRR-XI (Leucine Rich Repeat-XI) and annotate it with PAMP transcriptional data. For the gene tree in this case study, we include both the model plant *A. thaliana* (69) and *Solanum lycopersicum* (Tomato) (70), a major commercial crop where a comparable PAMP induction dataset is not available. The BAT pipeline produced a gene tree (**Fig. 2a**) that is consistent with previous work on *A. thaliana* RLK sub-family LRR-XI (71–73). Many of the functionally characterized developmental receptors in the tree show no change in expression following PAMP treatment. By contrast, HAESA-Like 3 (AtHSL3), the receptor for the immune peptide-hormone CTNIP4, shows strong PAMP induced upregulation in each treatment (**Fig. 2b**) (67). Notably, the PAMP data in Bjornson *et al*. was key to the discovery that *At*HSL3 is the receptor for the CTNIP4 immune peptide hormone (74), highlighting the value of integrating experimental data into the BAT analysis.

**Figure 2:**
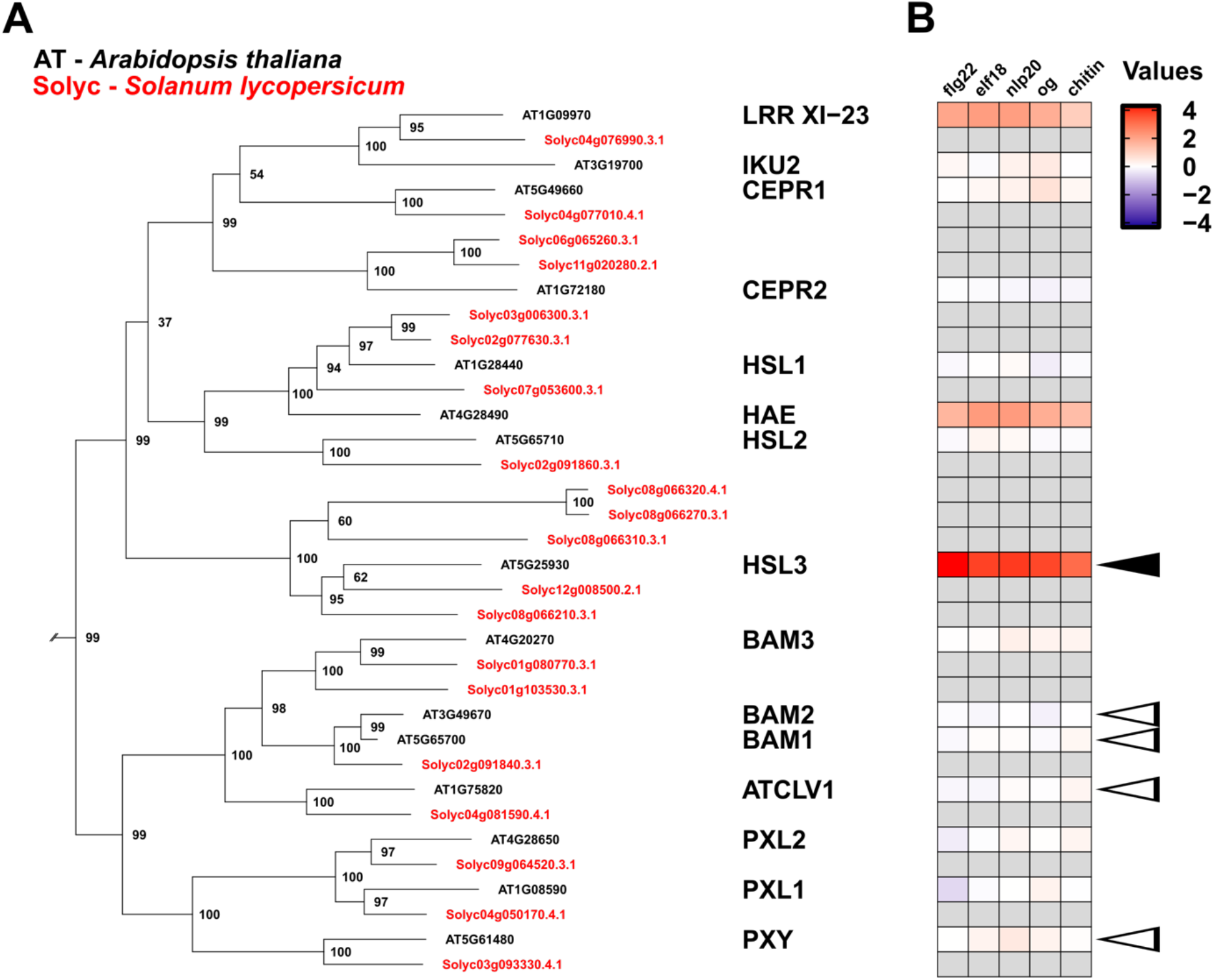
Annotation of a plant RLK-LRR subfamily XI gene tree subclade with RNA-seq data from *Bjornson* et al. 2021. BAT used the INTERPRO-Pfam identified kinase domain (amino acids 690-961) of *A. thaliana* transcript AT5G25930.1 (AtHSL3), as an option (-aa) BLAST query against the *A. thaliana* and *S. lycopersicum* genomes. See full query in **Supplemental Table 1**. Figure was adapted from BAT pipeline output. (**A**) The initial BAT tree was pruned to display only the HSL3/CLV1 clade (clade support = 0.61). Gene IDs and enlarged gene symbols colored according to species. (**B**) Heat map of log2-fold change in gene expression following treatment with the indicated PAMP (columns). Grey heat map cells indicate no data. “Values” is the default heatmap legend. Solid arrow highlights immune receptor AtHSL3, outlined arrows indicate characterized developmental receptors (AtCLV1, AtBAM1/2, AtPXY).

Adding additional genomes for gene tree exploration is simple with the BAT pipeline (See Methods). Here, the addition of tomato to the BAT gene tree highlights Solyc08g066210.3.1 and Solyc12g008500.2.1 as candidate CTNIP receptors based on orthology to AtHSL3. It further identifies a potentially tomato-specific sister clade of 3 HSL3-like homologs in tomato, which could diverge functionally from HSL3. While confirmation of these hypotheses requires experimental data, BAT analysis provides a foundation for these questions.

#### 2. BAT Facilitated Protein Domain Annotation of Gene Trees

Annotating protein domain architecture can further aid the analysis of gene families. To facilitate this analysis, BAT annotation can overlay HMMscan (75) identified domains on an alignment visualization in line with the gene tree. One example where domain architecture evolution is known to affect gene function is in the expanded plant immune NOD-like Receptor (NLR) family (76, 77). Many plant NLRs contain Integrated Domains (IDs), which facilitate interaction with pathogen proteins by mimicking the true host targets (such as transcription factors) (78). Plant NLRs genes are often encoded at tandemly duplicated loci that include nonfunctional paralogs (76, 79). Thus, identifying potential pseudo-genes and functional genes with novel domain architectures typically requires manual curation.

To demonstrate domain architecture annotation, we show the BAT gene tree of a rapidly evolving NLR subfamily. The *A. thaliana* gene Resistance to *Ralstonia solanacearum* 1 (RRS1) is a plant NLR immune receptor that recognizes the pathogen protein AvrRps4. RRS1 is a Toll/interleukin-1 Receptor (TIR) type NLR, characterized by the presence of an N-terminal TIR domain, a Nucleotide Binding-Apoptotic protease-activating factor-1 Resistance protein and *Caenorhabditis elegans* death-4 (NB-ARC) domain, and a C-terminal LRR domain (79). *At*RRS1 also has a WRKY transcription factor integrated domain which binds AvrRps4 (80, 81). RRS1 copy numbers vary between related species (76, 82).

BAT analysis of this gene family shows that genes outside the RRS1/b clade lack the WRKY domain, including the closest homologs in related plant species *Brassica rapa* and *Eutrema salsugineum* (**Fig. 3a-b**). Within the RRS1 clade, three of five genes from the congeneric species *Arabidopsis lyrata* also lack the WRKY domain, one of which is a possible pseudo gene that lacks canonical NLR domain architecture (**Fig. 3b**). BAT analysis suggests that only *At*RRS1/1b and *A. lyrata* genes AL8G14230 and AL8G14370 are candidate functional AvrRps4 receptors, and that the WRKY ID fusion is lineage specific. Further exploration of *A. thaliana* and *A. lyrata* homolog variation is facilitated by additional BAT analyses including the default multiple sequence alignment file where genes are ordered according to the tree (ggtree) outputs. This example demonstrates the utility of BAT for analyzing functionally relevant protein domain variation within a gene family.

**Figure 3:**
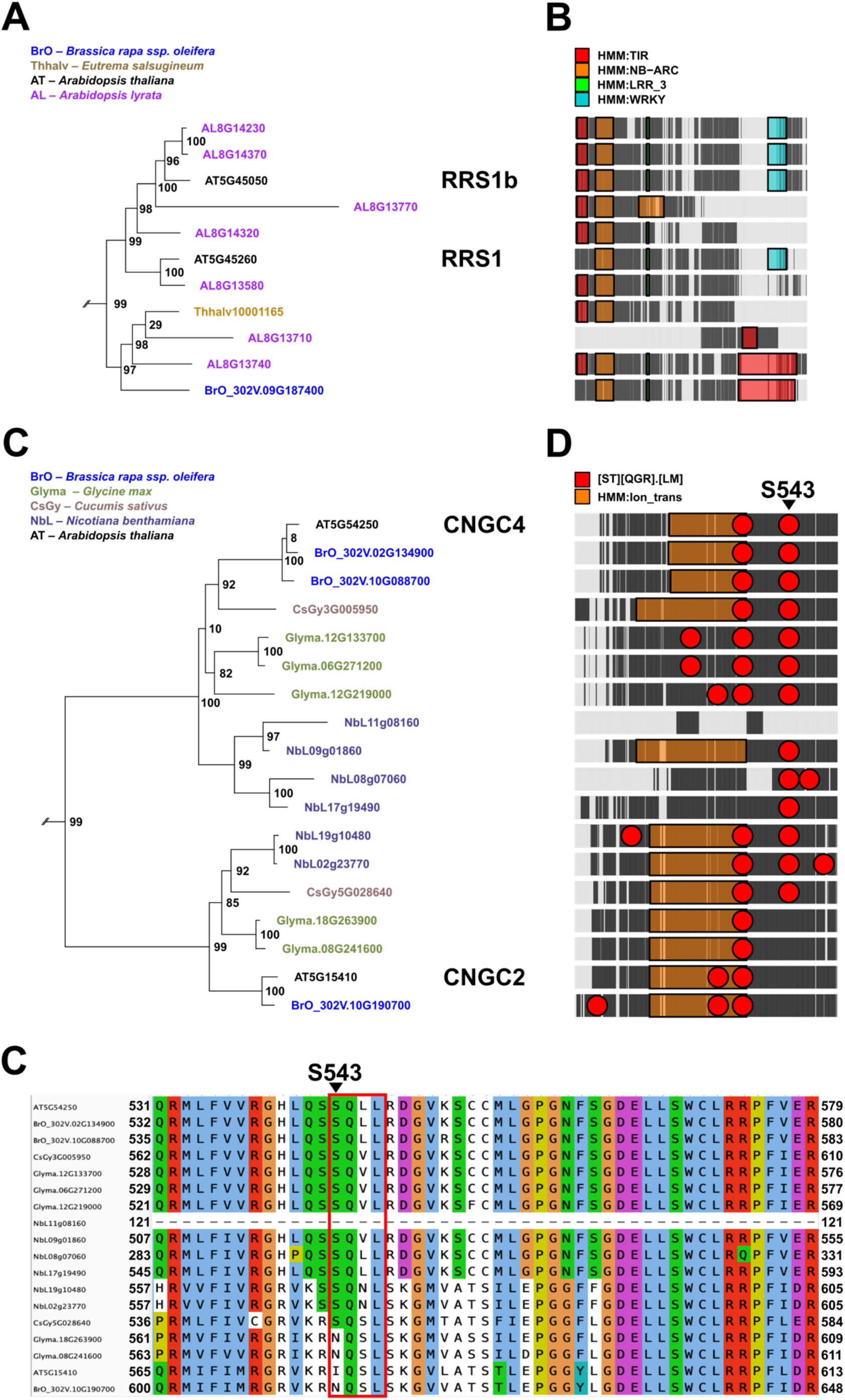
Annotation of plant NLR immune receptors and CNGC calcium ion channels with HMMscan protein domains and amino acid motifs. Gene IDs and enlarged gene symbols colored according to species. In alignment (MSA) diagrams, black blocks signify aligned sequence, grey spaces show alignment gaps, overlayed colored blocks correspond to indicated protein domains, and circles to targeted amino acid motifs. Pannels were adapted from BAT pipeline output. See full queries in **Supplemental Table 1**. (**A**) BAT used *A. thaliana* transcript AT5G45260.1 (AtRRS1) as a BLAST query against the *A. thaliana, A. lyrata, B. rapa*, and *E. salsugineum* genomes, pulling 50 hits in each search. The initial BAT tree was pruned using visualize_tree.r to display only the RRS1/1b clade (support = 99). (**B**) TIR, NB-ARC, LRR, and WRKY domains overlayed on RRS1 MSA visualization, aligned to tree (**C**) BAT used *A. thaliana* transcript AT5G54250.1 (AtCNGC4) as a BLAST query against the *A. thaliana, B. rapa, N. benthamiana, G. max*, and *C. sativus* genomes, pulling 50 hits in each search. The initial BAT tree was pruned to display only the CNGC2/4 clade (support = 86). (**D**) Ion Transport domains and motif [ST][QGR].[LM] annotation overlayed on CNGC2/4 MSA diagram, aligned to tree (**E**) BAT produced MSA corresponding to the gene tree in panel C.

#### 3. Motif Annotation of Gene Tree MSAs

Annotation of amino acid motifs is another routine task in gene family analysis. The BAT motif option enables the mapping of a specified amino acid motif onto the alignment visualization. To demonstrate the motif annotation feature, we highlight putative phosphorylation target sites in plant immune signaling. The amino acid motif SxxL (Serine followed by any two amino acids, followed by Leucine), is an established phosphorylation target of the plant Receptor Like Cytoplasmic Kinase (RLCK) Botrytis Induced Kinase1 (*At*BIK1) (83–86). *At*BIK1 is a conserved plant immune signaling kinase that links cell surface receptors to downstream potentiators of immunity including Cyclic Nucleotide Gated Channels (CNGCs) (85, 87). A heterodimer of *A. thaliana* CNGC2 and CNGC4 is responsible for Ca2+ influx during the early plant immune response and activates when *At*CNGC4 is phosphorylated by *At*BIK1 at S543. *At*CNGC2 by contrast, is not phosphorylated by *At*BIK1, and lacks phospho-site characterized in *At*CNGC4 (85).

To demonstrate BAT amino acid motif annotation, we annotated the *At*CNGC4-S543 phosphorylation site across flowering plant lineages using BAT (**Fig. 3c-e**). The gene tree in **Fig. 3c** supports a CNGC2/4 clade comprised of CNGC2, CNGC4, and their orthologs. The *At*CNGC4 S543 site is highlighted in the MSA diagram (**Fig. 3d**) and broadly conserved among *At*CNGC4 in other species. Interestingly, while Serine is absent at the equivalent position in *At*CNGC2 consistent with the literature, CNGC2 orthologs in *Cucumis sativus* (Cucumber) and *Nicotiana benthamiana*, have an intact SxxL (**Fig. 3d-e**). The conservation of this site suggests that it is ancestral within CNGC2/4s gene family. Potential loss-of-phosphorylation in the *At*CNGC2 lineage suggests that regulation of this homolog has evolved to be dispensable, with implications for the ancestral and derived mode of homo- and hetero-dimerization.

## DISCUSSION

Gene trees are critical for inferring gene function, especially for uncharacterized gene homologs in model and non-model organisms. Here we introduce BAT, a new bioinformatic pipeline for customizable gene tree homology search, construction, and annotation. Existing ortholog detection pipelines are well established at the genome scale (2, 3, 15–20). However, large-scale, all-against-all comparisons for pre-compiled ortholog databases are not appropriate for non-model organisms with custom or unpublished genome assemblies. Additionally, fine-scale phylogenetic analysis is often critical for rapidly evolving gene families where 1:1 orthologs are rare. Finally, this pipeline addresses the current lack of tools that combine comprehensive gene tree construction with automated data annotation and is a powerful inferential tool for gene discovery and gene family exploration.

The BAT pipeline generates trees for individual gene families and is complementary to genome-wide ortholog mining approaches. BAT’s bottom-up design offers several practical advantages for gene family characterization. First, researchers retain full control over taxon sampling, allowing genomes of interest, such as draft assemblies, to be incorporated simply by adding a new genome (FASTA) file. Second, BAT focuses on specific genes as BLAST queries to constrain the analysis to a manageable set of closely related homologs. Indeed, the large discrepancies in global orthogroup number between different approaches suggests great complexity with many gene families, and a need for fine-scale tree building for families of interest (3). Third, the BAT pipeline supports manual refinement which is especially useful in rapidly evolving gene families that often lack direct orthology. Users can adjust query ranges, swap aligners and/or tree-building methods, re-root trees, and extract subclades to interrogate complex 1:many and many:many gene family relationships.

Where re-visualization of outputs does not require new analysis, BAT allows iterative editing of tree display features without re-running the complete analysis. This approach complements genome-wide orthology resources by enabling fine-scale, hypothesis-driven phylogenetic analysis tailored to the specific gene family and species set under investigation. For example, BAT recovers known gene families including mammalian NAIPs and plant FLS2 orthologs (Supplemental Fig. 1, Supplemental Fig. 2). Within each gene family, BAT generated phylogenies can detect lineage-specific gene duplications (murid NAIPs) and losses (cucurbit FLS2). These events may be obscured in simple one-to-one ortholog tables or precomputed pairwise orthology summaries. By contrast, focused BAT trees allow users to inspect the local gene-family topology, sampling depth, annotation structure, and support values directly. With BAT, relationships relative to characterized genes from genetic model organisms are determined by the user, can be confirmed with different search, alignment, and tree building options, and further analyzed at the sequence level. BAT outputs include tree and sequence files which can be combined with methods such as NOTUNG or TreeRecs for species-gene tree reconciliation (88, 89), and Possvm for fine-scale ortholog determination (90).

Automated pipelines can be dangerous; a poorly built tree will lead to false conclusions. The BLAST, alignment, and tree building steps of BAT are the foundation for later analysis, and each step still requires decision making by the user. While all key packages implemented in the BAT pipeline are previously benchmarked (6–8, 12, 13), interpretations of BAT-generated gene trees are only valid if the user fully understands the implications of choices made in the pipeline (i.e., at the point of specifying pipeline parameters via terminal commands). The examples above were generated with BAT searches that gather an abundance of BLAST hits (option -n 50). Larger runs using hundreds of hits are advisable before drawing strong phylogenetic conclusions. Relatedly, the option -hdr parses fasta file headers for appropriate portions of sequence descriptions to identify and manage sequences. For example, a user can specify tree construction using transcript vs gene labels to either retain or condense splice isoforms. It is important to consider that the header parsing and deduplication step takes place after the BLAST search. If the query blasts primary-transcript-only files and isoform containing files in the same search, the effect is to recover fewer genes from FASTAs with isoforms. To address this, the (-n) parameter can be specified independently for each genome in the query to account for species-specific expansions or recent whole genome duplications.

For alignment and tree building options, different combinations are possible. We recommend starting with a lightweight analysis (Clustal Omega (7) + FastTree (13)), adding additional genomes, analysis, and annotation in subsequent runs. BAT incorporates MAFFT (8) as an alternative alignment tool which allows users to specify the alignment strategy. MAFFT’s iterative alignment strategy may provide greater accuracy for some datasets compared with Clustal Omega’s linear progressive strategy. However, this comes at a computational cost for large datasets. RAxML-NG (12) is available in BAT as an alternative tree construction tool. RAxML-NG produces rigorous maximum likelihood gene trees for publication. Like MAFFT, RAxML-NG scales non-linearly and is similarly computationally expensive for large datasets. BAT delegates alignment and tree inference to widely used, previously benchmarked tools, while adding reproducible local homolog retrieval and integrated downstream annotation.

A strength of BAT is its ability to combine gene trees with experimental data to identify genes of interest across species. The BAT-generated *A. thaliana* gene tree combined with the transcriptomic Bjornson *et al*. (68) dataset heatmap demonstrates how this analysis can lead to gene discovery (Fig 2). Specifically, it highlights HSL3 as transcriptionally responsive to PAMPs when compared to related receptors, leading to identification of candidate genes for further study. This specific observation originally supported the discovery that HSL3 is the CTNIP receptor. Additionally, BAT annotations facilitate the extrapolation of functional data to orthologous genes. The gene tree in Case Study 1 defines orthologous relationships between *A. thaliana* genes with transcriptomic data, and tomato orthologs where no data is available.

Although functional conservation is not guaranteed across species, cross-referencing the PAMP induced expression of *A. thaliana* genes with tomato homologs provides a powerful basis for identifying tomato genes of interest in the absence of tomato specific data. In order to facilitate general BAT use in plants, we provide a default expression data subset for *A. thaliana* sourced from the Klepikova expression atlas as a BAT default (91).

Structural annotation of gene tree sequences provides further context for putative gene functional and evolutionary inference. BAT annotation of the *At*RRS1 NLR gene tree with both canonical and noncanonical NLR protein domains illustrates the effectiveness of this strategy for gene family exploration. Annotation with canonical NLR domains rapidly identified one *A. lyrata* gene within the RRS1/b clade that lacks core machinery for NLR function. Conversely, annotation with the non-canonical but AvrRps4 sensor function critical WRKY domain highlights two putative *A. lyrata* receptors while excluding two others and confirming lineage specificity. Intriguingly, the alignment visualization (**Fig. 3b**) shows that one of these paralogs (AL8G13580) shares many blocks of aligned sequence with the WRKY domain of *At*RRS1 suggesting partial degradation of WRKY. If this RRS1 homolog indeed lacks AvrRps4 sensor capability, analysis of its C-terminal domain could identify residues critical for WRKY-AvrRps4 interaction.

BAT gene tree annotation of amino acid motifs reveals similarly important patterns. In *A. thaliana*, only CNGC4 conserves the BIK1 phospho-motif S543 where it is necessary for CNGC2/4 heterodimer activation. However, analysis of CNGC2/4s shows that the S543 SxxL motif is ancestral in the CNGC2/4 clade and multiple species retain the site in both genes. This analysis raises the possibility that BIK1 phosphorylates CNGC2 in some species, suggesting possible differences in CNGC2/4 channel phosphorylation and opening during immunity. However, all orthology-based functional inferences must be validated experimentally. Nonetheless, these case studies highlight BATs power as an exploratory tool to infer protein functional characteristics.

User computational power is a limiting factor for BAT. However, BAT is not inherently a heavyweight pipeline. Both default alignment and tree construction options Clustal Omega and FastTree work well when datasets are large or computational power is limited. BAT analysis for all case studies presented here were performed on a MacBook Pro in seconds to minutes with these options. Increasing genome count, BLAST depth, or utilizing alternate aligner MAFFT and tree builder RAxML-NG can increase breadth and analytical rigor at computational cost. To address this, BAT automatically parallelizes BLAST searching and utilizes computational threads for alignment and tree construction to reduce analysis time. Consequently, BAT scales effectively for computationally expensive queries when computational power is available.

In summary, BLAST-Align-Tree uniquely integrates experimental and sequence structure data with phylogenetic analysis in a single pipeline, enhancing homolog discovery and facilitating functional hypothesis generation.

## Implementation Note

BAT is implemented in Python and uses R for visualization. It is freely available under an open-source license from the steinbrennerlab/blast-align-tree GitHub repository (https://github.com/steinbrennerlab/blast-align-tree) and as a pip-installable package (pip install blast-align-tree). BAT runs from the command line and includes a Tkinter GUI (bat-genome-selector) for assembling runs. Dependencies are managed through platform-specific conda environment files compatible with conda, mamba, or micromamba. The Linux environment is a one-step install. MacOS users additionally install R and R dependencies from CRAN, and Windows users install the external CLI tools (e.g. BLAST) from vendor binaries. Alternatively, Windows users can run the Linux environment under WSL2. Complete installation instructions, tutorials, and worked examples are provided in the repository.

## Acknowledgements

This work was supported by National Institutes of Health (NIH) and National Science Foundation (NSF) awards to ADS (5R35GM151272 and 2139986 respectively).

