## Supplemental Figures and Tables for "BAT: an integrated pipeline for gene tree construction, annotation, and functional inference"

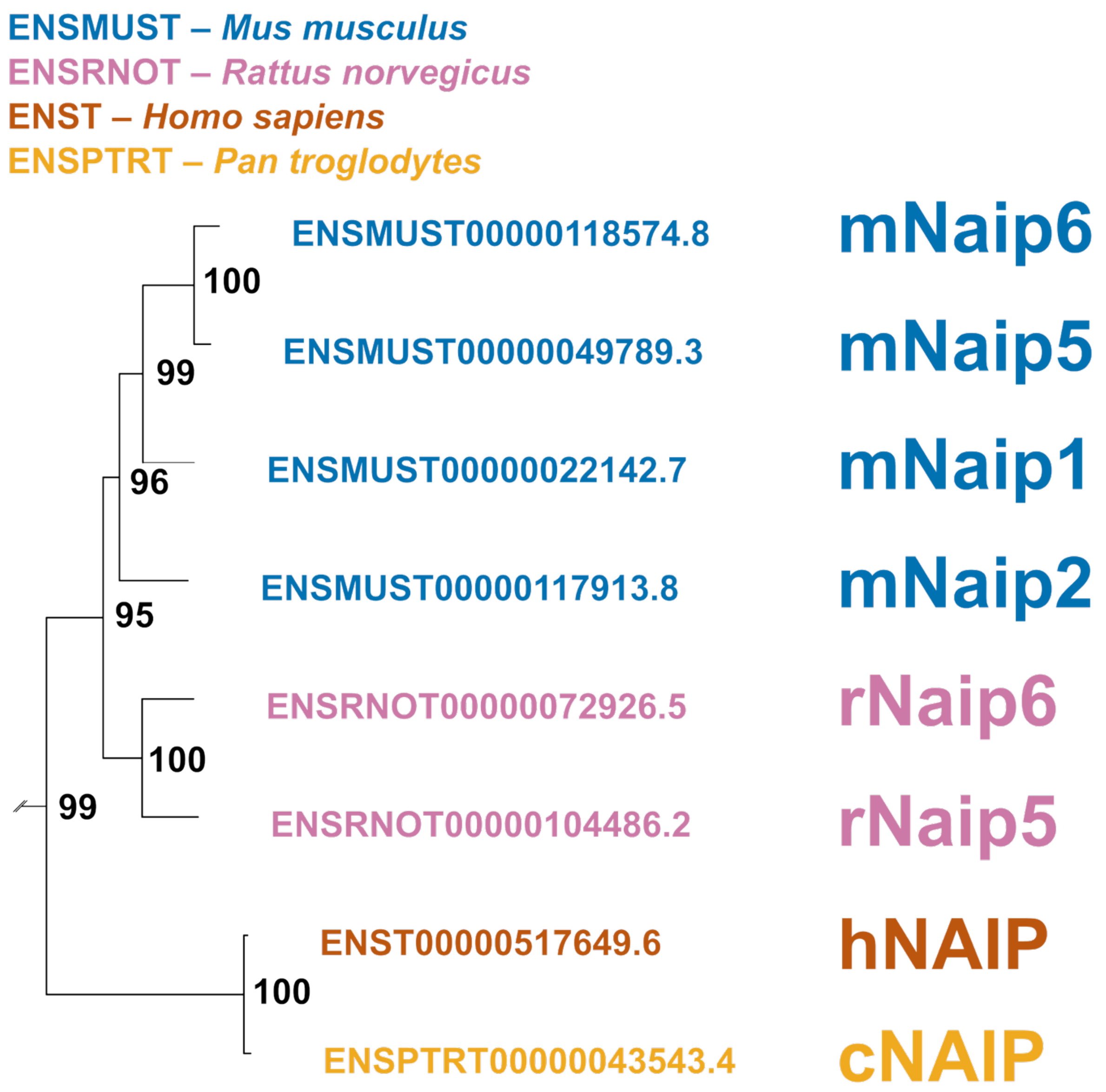


**Supplemental Figure 1:** BAT gene tree analysis of NAIPs from four mammalian species. BAT used Human NAIP (ENST00000517649.6) as a BLAST query against *Homo sapiens, Pan troglodytes, Mus musculus,* and *Rattus norvegicus* genomes, pulling 30 hits in each search. See full query in **Supplemental Table 1**. Figure was adapted from BAT pipeline output and pruned to display only the NAIP clade (clade support = 82). Gene IDs and gene symbols colored according to species.

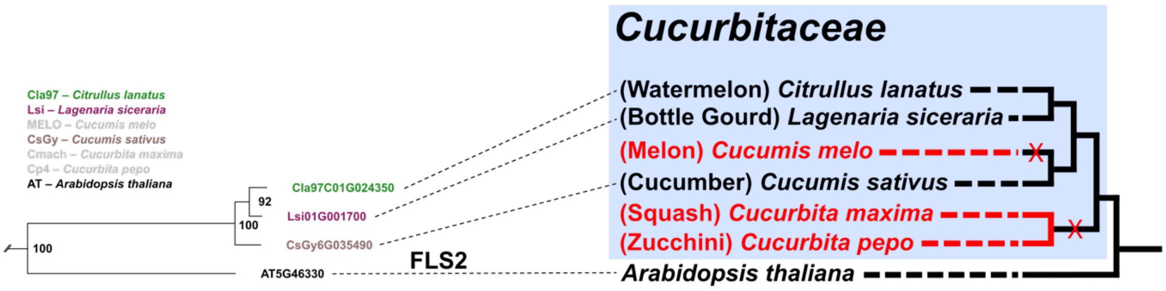


**Supplemental Figure 2:** BAT analysis of plant FLS2 orthologs in Cucurbitaceae manually aligned to species cladogram. BAT blasted *A. thaliana* FLS2 against *A. thaliana, C. lanatus, L. siceraria, C. melo, C. sativus, C. maximum*, and *C. pepo*, pulling 50 hits in each search. See full query in **Supplemental Table 1**. Gene tree was adapted from BAT pipeline output and pruned to display only the FLS2 clade (clade support = 0.98). Gene IDs and gene symbols colored according to species. Cladogram tips and branches colored by loss of FLS2 ortholog (red). Red "X"s indicates inferred lineage-specific loss of FLS2. Blue box indicates species within Cucurbitaceae (*A. thaliana* outgroup).

**Supplemental Tabe 1:** BLAST-Align-Tree Terminal Commands for Figures

| Figure | Query Type | Terminal |
|---|---|---|
| S1 | P | Blast-align-tree -q ENST00000517649.6 -qdbs human_cds_canonical.fa -n 30 30 30 30 -dbs human_cds_canonical.fa mouse_cds_canonical.fa rat_cds_canonical.fa chimp_cds_canonical.fa -hdr '>' '>' '>' '>' |
| S1 | V | Rscript "/[FILE PATH]/blast_align_tree/data/visualize_tree.r" -e ENST00000517649.6 -b ENST00000517649.6 --subdir runs/20260310_1710 -k 1 -l 0 |
| S2 | P | blast-align-tree -q AT5G46330.1 -qdbs TAIR10cds.fa -dbs CM4.0_transcripts.fasta Cmaxima_v1.1.cds.fa Cpepo_v4.1.cds.fa Gy14_v2.cds.fa TAIR10cds.fa USVL1VR-Ls_CDS_v1.fa watermelon_97103_v2.fa -hdr id id id id gene: id id -hdr_sfx .1 .1 .1 .1 none .1 .1 -n 50 50 50 50 50 50 50 |
| S2 | V | Rscript "/[FILE PATH]/blast_align_tree/data/visualize_tree.r" -e AT5G46330.1 --subdir runs/20260426_1606 -k 1 -l 0 -m 2 -b fls2clade -n 381 --genome_colors 'TAIR10cds.fa=black, Gy14_v2.cds.fa=rosybrown4, watermelon_97103_v2.fa=forestgreen, USVL1VR-Ls_CDS_v1.fa=maroon4' |
| 2 | P | blast-align-tree -q AT5G25930.1 -qdbs TAIR10cds.fa -dbs Solyc_ITAG4.0_CDS.fa TAIR10cds.fa -hdr id gene: -n 50 50 -aa 690 961 --datasets datasets/steinbrenner_lab/bjornsen_pamps_l2FC.txt |
| 2 | V | Rscript "/[FILE PATH]/blast_align_tree/data/visualize_tree.r" -e AT5G25930.1 -b HSL3clade --subdir runs/20260427_1631 -n 96 -k 1 -l 0 -m 2 --datasets datasets/steinbrenner_lab/bjornsen_pamps_l2FC.txt --genome_colors 'TAIR10cds.fa=black,Solyc_ITAG4.0_CDS.fa=red' |
| 3a-b | P | blast-align-tree -q AT5G45260.1 -qdbs TAIR10cds.fa -dbs Alyrata_384_v2.1.cds_primaryTranscriptOnly.fa BrapaO_302V_711_v1.1.cds.fa Esalsugineum_173_v1.0.cds.fa TAIR10cds.fa -hdr locus= locus= id gene: -hdr_sfx none none m none -n 50 50 50 50 --hmm WRKY.hmm NB-ARC.hmm LRR_3.hmm TIR.hmm |
| 3a-b | V | Rscript "/[FILE PATH]/blast_align_tree/data/visualize_tree.r" -e AT5G45260.1 --subdir runs/20260426_1117 -k 1 -l 0 -m 2 -b rrs1clade -n 140 --genome_colors 'TAIR10cds.fa=black, Alyrata_384_v2.1.cds_primaryTranscriptOnly.fa=purple, BrapaO_302V_711_v1.1.cds.fa=blue, Esalsugineum_173_v1.0.cds.fa=darkgoldenrod' |
| 3c-e | P | blast-align-tree -q AT5G54250.1 -qdbs TAIR10cds.fa -dbs BrapaO_302V_711_v1.1.cds.fa Gmax_880_Wm82.a6.v1.cds.fa Gy14_v2.cds.fa NbLab360.v103.gff3.CDS.fa TAIR10cds.fa -hdr locus= locus= id id gene: -hdr_sfx none none .1 .1 none -n 50 50 50 50 50 --motif '[ST][QGR].[LM]' --hmm Ion_Transport.hmm |
| 3c-e | V | Rscript "/[FILE PATH]/blast_align_tree/data/visualize_tree.r" -e AT5G54250.1 --subdir runs/20260426_1206 -k 1 -l 0 -m 2 -b cngc24clade -n 169 --genome_colors 'TAIR10cds.fa=black, Gmax_880_Wm82.a6.v1.cds.fa=darkolivegreen4, NbLab360.v103.gff3.CDS.fa=darkslateblue, BrapaO_302V_711_v1.1.cds.fa=blue, Gy14_v2.cds.fa=rosybrown4' |

**Supplemental Tabe 2:** BLAST-Align-Tree vs eFP Browser Klepikova Atlas Select Genes

| **AT2G32680** | **RLP23** | | |
|---|---|---|---|
| Tissue | BAT log2 | eFP RPKM (avg) | log2(RPKM+1) |
| Root | 2.04 | 0.067 | 0.09 |
| Carpel | 2.46 | 0.1 | 0.14 |
| Petal | 0 | 0 | 0 |
| Internode | 4.19 | 0.28 | 0.36 |
| Dry seed | 2.1 | 0.025 | 0.04 |
| Mature leaf | 7.43 | 2.539 | 1.82 |
| | | PEARSON: | 0.922 |

| **AT4G31380** | **FLP1** | | |
|---|---|---|---|
| Tissue | BAT log2 | eFP RPKM (avg) | log2(RPKM+1) |
| Root | 0 | 0 | 0 |
| Carpel | 3.84 | 1.768 | 1.47 |
| Petal | 5.61 | 5.294 | 2.65 |
| Internode | 0 | 0 | 0 |
| Dry seed | 0 | 0 | 0 |
| Mature leaf | 7.38 | 18.459 | 4.28 |
| | | PEARSON: | 0.984 |

| **AT4G33430** | **BAK1** | | |
|---|---|---|---|
| Tissue | BAT log2 | eFP RPKM (avg) | log2(RPKM+1) |
| Root | 10.86 | 21.03 | 4.46 |
| Carpel | 10.1 | 12.841 | 3.79 |
| Petal | 9.64 | 7.502 | 3.09 |
| Internode | 10.34 | 13.682 | 3.88 |
| Dry seed | 10.82 | 6.666 | 2.94 |
| Mature leaf | 10.39 | 12.853 | 3.79 |
| | | PEARSON: | 0.347 |

| **AT3G27810** | **MYB21** | | |
|---|---|---|---|
| Tissue | BAT log2 | eFP RPKM (avg) | log2(RPKM+1) |
| Root | 0 | 0 | 0 |
| Carpel | 11.24 | 27.307 | 4.82 |
| Petal | 13.23 | 86.044 | 6.44 |
| Internode | 2.03 | 0.031 | 0.04 |
| Dry seed | 1.08 | 0.008 | 0.01 |
| Mature leaf | 1.85 | 0.024 | 0.03 |
| | | PEARSON: | 0.991 |

| **AT2G02760** | **UBC2** | | |
|---|---|---|---|
| Tissue | BAT log2 | eFP RPKM (avg) | log2(RPKM+1) |
| Root | 10.97 | 47.740 | 5.61 |
| Carpel | 10.45 | 33.510 | 5.11 |
| Petal | 11.24 | 46.671 | 5.58 |
| Internode | 11.58 | 65.366 | 6.05 |
| Dry seed | 11.82 | 27.676 | 4.84 |
| Mature leaf | 11.98 | 79.853 | 6.34 |
| | | PEARSON: | 0.407 |
